# Extensive memory testing improves prediction of progression to MCI in late middle age

**DOI:** 10.1101/584193

**Authors:** Daniel E. Gustavson, Jeremy A. Elman, Mark Sanderson-Cimino, Carol E. Franz, Matthew S. Panizzon, Amy J. Jak, Chandra A. Reynolds, Michael C. Neale, Michael J. Lyons, William S. Kremen

## Abstract

**INTRODUCTION:** Predicting risk for Alzheimer’s disease when most people are likely still biomarker negative would aid earlier identification. We hypothesized that combining multiple memory tests and scores in middle-aged adults would provide useful, and non-invasive, prediction of 6-year progression to MCI.

**METHODS:** We examined 849 men who were cognitively normal at baseline (mean age=55.69±2.45).

**RESULTS:** California Verbal Learning Test learning trials was the best individual predictor of amnestic MCI (OR=4.75). A latent factor incorporating 7 measures across 3 memory tests provided much stronger prediction (OR=9.88). This compared favorably with biomarker-based prediction in a study of much older adults.

**DISCUSSION:** Neuropsychological tests are sensitive and early indicators of Alzheimer’s disease risk at an age when few individuals are likely to have yet become biomarker positive. Single best measures may appear time- and cost-effective, but 30 additional minutes of testing, and use of multiple scores within tests, provides substantially improved prediction

---

The pathogenesis of Alzheimer’s disease (AD) begins decades before the onset of dementia, so it is necessary to identify risk factors as early as possible[1-4]. The recent A/T/N (amyloid/tau/neurodegeneration) framework emphasizes biomarkers in an effort to improve early identification and move toward a biological diagnosis[5, 6]. Preclinical AD is stage 1 of the A/T/N framework staging, defined as being amyloid positive but still cognitively normal. However, being biomarker positive indicates that a significant amount of disease progression has already taken place. The ability to identify individuals at risk *before* they become biomarker positive would thus be of great potential value. It would also be useful to identify people who are likely to be at elevated risk before embarking on costly and invasive biomarker testing. This view echoes those of several research groups who have noted there is pressing need to identify tests that are non-invasive, low-cost, and can improve earlier identification of risk for AD[4, 7-9].

Episodic memory is an effective early predictor of progression to AD[10-15]. It should also be a good predictor of amnestic mild cognitive impairment (MCI). Memory is severely impaired in AD, and MCI diagnoses are in part based on impaired memory performance[16]. However, comprehensive neuropsychological assessment of memory (and other cognitive domains) is often lacking in longitudinal studies aimed at detecting which individuals are at the greatest risk for MCI/AD. It is also important to recognize that cognitively normal individuals are not a homogeneous group. Examining variability within the range of normal cognitive function may be useful for early prediction of MCI.

Efforts to improve longitudinal prediction of MCI or AD using cognitive measures have often focused on evaluating which measures provide the best prediction compared with other measures[15, 17, 18]. This approach also rests on an unspoken assumption that including the lesser predictors in the model may not improve (or may even hurt) predictive ability. However, we have shown that multiple memory tests (and scores within tests) have both common and unique genetic and environmental influences[19, 20], suggesting that the right combination of tests might enhance prediction. In the current study, we tested the hypothesis that combining multiple memory measures (within test and across multiple tests) provides stronger and more robust prediction than the single best measure, and that differences, even among relatively younger middle-aged adults, can be predictive of progression to MCI.

We evaluated our hypothesis in a community sample of middle-aged adults from the Vietnam Era Twin Study of Aging (VETSA) who were all cognitively normal at baseline. They completed multiple memory tests at mean ages 56 and 62. Examining these associations in midlife is important because improving treatment efficacy may depend on early intervention[7, 21], and cognitive abilities may already be subtly declining by the late 50s[20, 22-24]. Yet there is exceedingly little focus on prediction of MCI, particularly in adults this young. In these analyses, we compared two approaches to aggregating measures: *z*-score composites and factor scores. We expected that the factor score approach would have the strongest prediction because it weighs more strongly the measures that are the best indicators of the latent memory construct.

## Method

### Participants

Analyses were based on 849 individuals from the longitudinal Vietnam Era Twin Study of Aging (VETSA) project who were cognitively normal at wave 1, returned to complete the wave 2 assessment approximately 6 years later, and had data for all covariates. Participants were recruited randomly from a previous large-scale study of Vietnam Era Twin Registry participants[25]. All served in the United States military at some time between 1965 and 1975; nearly 80% did not serve in combat or in Vietnam[26, 27]. Participants are generally representative of American men in their age group with respect to health and lifestyle characteristics[28]. All participants provided informed consent and the study was approved by local Institutional Review Boards at the University of California, San Diego and Boston University.

Of the 1,237 individuals who completed the VETSA protocol at wave 1, 107 (8.6%) were excluded from the analysis of progression to MCI because they had MCI at wave 1. This left 1130 (91.4%) cognitively normal individuals, 906 (80.2%) of whom returned for wave 2. Of those 906, 57 were excluded for missing covariates. This left 849 (93.7% of 906 and 75.1% of the total CN individuals at wave 1 for the analyses of progression to MCI. To compute standardized memory scores used in the odds ratios for the primary analyses, we used data from all 1,237 individuals at wave 1, plus an additional 53 attrition replacements who completed the VETSA protocol for the first time during the wave 2 assessment but were in the age range of participants at wave 1 (total N=1290). All subjects with available data were used so that odds ratios should better reflect those that would be obtained from a large population sample.

### Episodic Memory Measures

Episodic memory was measured at both waves with the Logical Memory (LM) and Visual Reproductions (VR) subtests of the Wechsler Memory Scale-III[29], and the California Verbal Learning Test-II (CVLT)[30]. For LM and VR, we examined immediate recall and delayed recall measures. For the CVLT, we examined the short delay and long delay free recall measures, and total score for learning trials 1-5). In analyses involving MCI, all memory measures were *z*-scored and transformed so that odds ratios (ORs) reflect the increase in odds of MCI for every decrease of 1 *SD* in memory performance (in relation to the full sample of N=1290 at wave 1). These standardization procedures were also conducted for the aggregated memory measures described next.

In addition to examining the predictive ability of each memory measure alone we combined measures in two ways. First, we created *z*-score composites for measures within a given test (e.g., LM immediate and delayed recall), and comparable measures across tests (e.g., LM immediate recall, VR immediate recall, CVLT short delay free recall). We also created *z-* score composites that combined the 6 short/long delay conditions across all tests or all 7 memory measures (including CVLT learning trials).

Second, we created factor scores from latent memory variables. Latent factors were exported from structural equation models in MPlus version 7.2[31] based in part on those reported in earlier work from this sample (see supplement for more information)[19, 20]. These factor scores are similar to the *z*-score composites, but measures that have stronger factor loadings on the latent memory factor are weighted more heavily. Latent factors are displayed in Figure 1 and were also based on the full wave 1 sample (N=1289). Each model had good fit to the data based on standard structural equation metrics, including Root Mean Square Error of Approximation values < .06, and the Comparative Fit Index values > .950[32].

**Figure 1:**
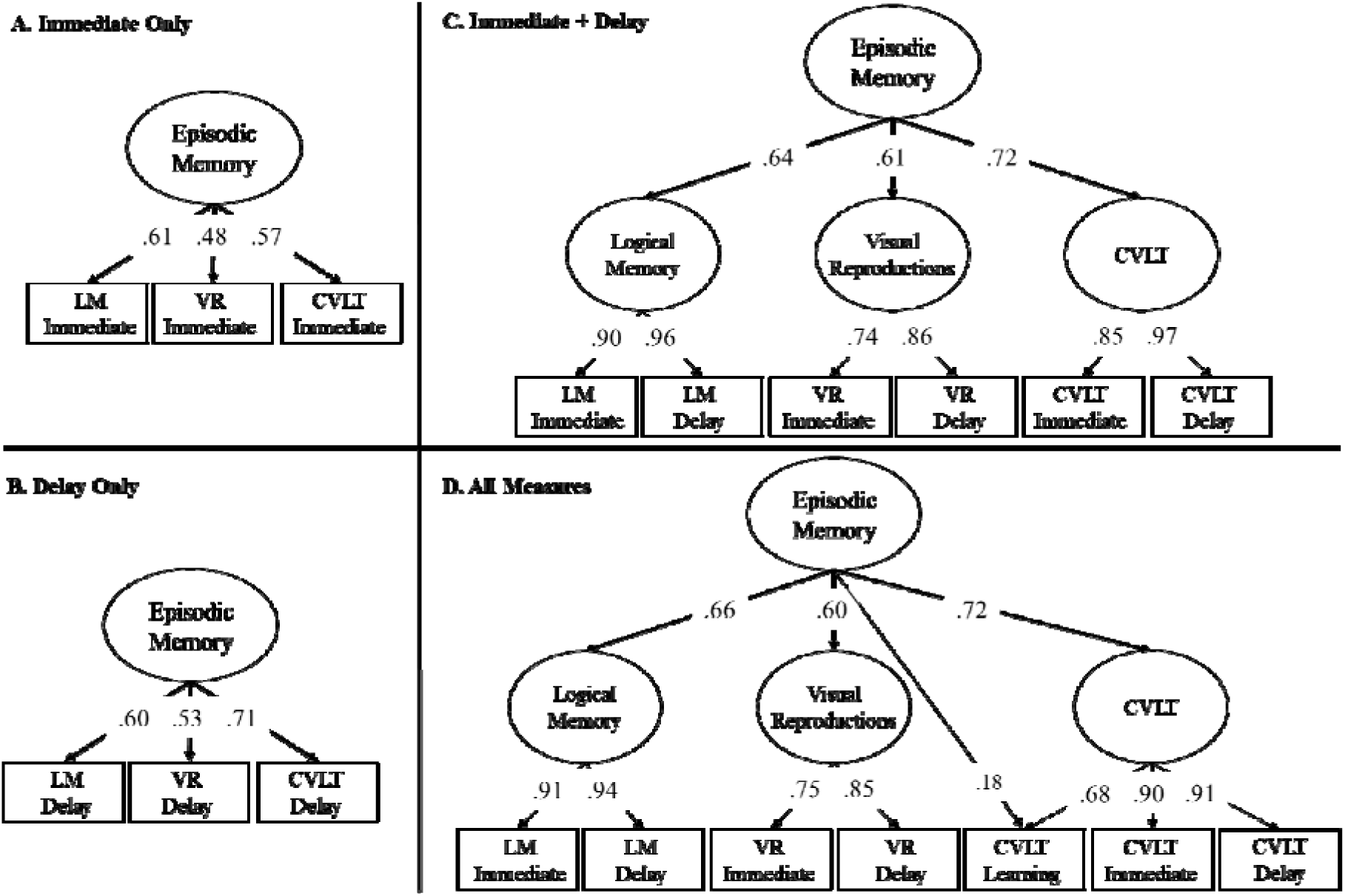
Structural equation models used to create factor scores for episodic memory at wave 1. In each model, variance explained in a given memory measure (rectangle) by latent factors (ovals) can be computed by squaring the factor loading on that factor. Factor scores for the highest-level Episodic Memory latent factor in each model were exported in Mplus and used as a continuous variable in separate logistic regression analyses involving MCI. All factor loadings were significant (*p* < .05) and all models fit the data well.

### Mild Cognitive Impairment Diagnoses

MCI was diagnosed using the Jak-Bondi approach[4, 16, 33]. Impairment in a cognitive domain was defined as having at least two tests >1.5 SDs below the age- and education-adjusted normative means after accounting for “premorbid” cognitive ability by adjusting neuropsychological scores for performance on a test of general cognitive ability that was taken at a mean age of 20 years[34]. The adjustment for age 20 cognitive ability ensures that the MCI diagnosis is capturing a decline in function rather than long-standing low ability. Most prior studies have used an impairment criterion on >1 SD, but they have had considerably older samples. The validity of the VETSA MCI diagnoses is supported in the present sample by evidence of reduced hippocampal volume in those diagnosed with amnestic MCI[35]. Higher AD polygenic risk scores were also associated with significantly increased odds of MCI in this sample[36], indicating that the MCI diagnosis is genetically-related to AD. MCI diagnoses at wave 2 were also based on measures that were adjusted to account for practice effects[37], leveraging data from attrition replacement subjects who completed the task battery for the first time at wave 2 (N=179), to estimate the increase in performance expected in returnees who completed the tests twice.

Because we were interested in transition to MCI, analyses included only individuals who were cognitively normal at wave 1 and had data for all covariates. Of the 849 returnees meeting this criterion, 45 (5.3%) progressed to amnestic MCI, and 41 (4.8%) progressed to non-amnestic MCI.

### Data Analysis

All statistical analyses were conducted using R version 3.5.1. Analyses involving MCI were conducted with mixed effects logistic regression using the lme4 package[38]. In these analyses, we controlled for wave 1 age, the time interval between assessments, education, race/ethnicity (white non-Hispanic vs. other), wave 1 diabetes (yes/no), wave 1 hypertension (yes/no), *APOE-*ε4 (ε4+ vs. ε4-), and wave 1 depression symptoms based on Center for Epidemiologic Studies–Depression scale[39]. Diabetes and hypertension status were based on whether the participant either (a) reported being diagnosed by a doctor, (b) reported they were currently taking medication for diabetes or high blood pressure, and/or (c) whether they had high blood pressure on the day of testing (hypertension only). Finally, twin pair ID was included as a random effect to account for the clustering of data within families. The lme4 package uses list-wise deletion with missing observations, and reports profile-based 95% confidence intervals (95% CIs).

The predictive utility of all models was assessed with average area under the curve (AUC) calculated from 4-fold cross-validation with 10 repeats. Random effects were not included in the cross-validated models due to difficulty obtaining model convergence with decreased sample size of MCI cases in each fold. However, we found that removing random effects from the full models resulted in decreased odds ratios of cognitive scores. Therefore, these AUC values may represent conservative estimates.

## Results

### Descriptive Statistics

There were no significant differences between cognitively normal and amnestic MCI groups in any demographic or clinical characteristics (Table 1). Compared to cognitively normal returnees, those diagnosed with non-amnestic MCI at wave 2 were older at baseline (*p*=.024) and a smaller proportion were white non-Hispanic (*p*=.031). ApoE4 was not a significant predictor, but it was in the expected direction for amnestic MCI (33% in amnestic MCI vs. 30% in controls). There is, however, evidence that ApoE effects are not as strong in men as in women and may be less prominent in our relatively young sample[40, 41]. Descriptive statistics for memory measures at wave 1 are displayed in Table 2.

**Table 1.** Demographic and Clinical Characteristics and Covariates Included in Analyses Involving Mild Cognitive Impairment (MCI)

| Demographic Variable | Remained Cognitively Normal (N = 763) | | Progressed to Amnestic MCI (N = 45) | | Progressed to Non-Amnestic MCI (N = 41) | | $p$ (CN vs. aMCI) | $p$ (CN vs. nMCI) |
| --- | --- | --- | --- | --- | --- | --- | --- | --- |
| | $M$ | $SD$ | $M$ | $SD$ | $M$ | $SD$ | | |
| Lifetime Education (years) | 14.03 | 2.16 | 13.58 | 1.75 | 13.49 | 1.83 | 0.191 | 0.098 |
| Age (at wave 1) | 55.69 | 2.45 | 56.37 | 2.48 | 56.68 | 2.41 | 0.102 | **0.024** |
| Age Interval (wave 2 - wave 1) | 5.75 | 0.70 | 5.76 | 0.67 | 5.65 | 0.54 | 0.907 | 0.400 |
| Depression Symptoms (at wave 1) | 7.69 | 7.14 | 7.32 | 7.03 | 9.71 | 9.66 | 0.858 | 0.152 |
| Ethnicity (% white non-Hispanic) | 91.34 | - | 84.44 | - | 80.49 | - | 0.119 | **0.031** |
| ApoE status (% ε4 positive) | 30.80 | - | 33.33 | - | 21.95 | - | 0.480 | 0.193 |
| Diabetes (% yes at wave 1) | 10.35 | - | 11.11 | - | 17.07 | - | 0.868 | 0.267 |
| Hypertension (% yes at wave 1) | 58.32 | - | 64.44 | - | 73.17 | - | 0.230 | 0.069 |

*Note:* The final two columns display the $p$ value for comparisons between individuals who remained cognitively normal (N=763) and either progressed to amnestic MCI (aMCI; N=45) or non-amnestic MCI (nMCI; N=41), controlling for clustering of data within families. Significant group differences are displayed in bold ( $p < .05$ ). Lifetime education was the number of years of school completed. Depression symptoms were measured with the Center for Epidemiologic Studies – Depression Scale [39], with scores above 15 indicating risk for clinical depression. Rate of ApoE ε4 positivity across the entire sample (30.5%) is similar to 28.9% in the UK Biobank, the largest population sample of ApoE prevalence to date (n=326,535) [50].

**Table 2.** Descriptive Statistics for Episodic Memory Measures at Baseline (Mean Age 56)

| Memory Variable | N | M | SD | Range | Skewness | Kurtosis |
| --- | --- | --- | --- | --- | --- | --- |
| *Logical Memory* |  |  |  |  |  |  |
| Immediate Recall | 1281 | 23.47 | 6.16 | 4, 44 | -0.04 | -0.10 |
| Delayed Recall | 1279 | 20.01 | 6.63 | 0, 41 | -0.10 | -0.13 |
| *Visual Reproductions* |  |  |  |  |  |  |
| Immediate Recall | 1284 | 78.24 | 12.41 | 21, 103 | -0.58 | 0.48 |
| Delayed Recall | 1283 | 54.75 | 19.51 | 0, 100 | -0.15 | -0.44 |
| *California Verbal Learning Test* |  |  |  |  |  |  |
| Learning Trials | 1270 | 42.84 | 8.51 | 18, 74 | 0.09 | -0.06 |
| Short Delay Free Recall | 1270 | 8.64 | 2.74 | 1, 16 | 0.07 | -0.21 |
| Long Delay Free Recall | 1269 | 9.06 | 2.89 | 0, 16 | 0.00 | -0.30 |

*Note:* In all analyses involving MCI, dependent measures were standardized and reverse scored so that odds ratios reflect increase in risk of MCI at lower levels of cognitive ability. Shown here are descriptive statistics for all individuals who completed the wave 1 protocol (used to standardize data and create z-score composites and latent factor scores). Distributional characteristics remained acceptable for the subset of 849 individuals in the analyses involving progression to MCI (range skewness = -.45 to .22; range kurtosis = -.36 to .05; see Table S1).

### Six-Year Prediction of Progression to Mild Cognitive Impairment

The results of the primary analyses are displayed in Table 3. Each cell displays an odds ratio from a separate longitudinal logistic regression in which a memory measure predicts progression to amnestic MCI (Table 3a) or non-amnestic MCI (Table 3b) 6 years later. As shown in Table 3a, each of the 7 individual memory measures at wave 1 significantly predicted progression to amnestic MCI at wave 2. The learning trials measure of the CVLT provided the strongest individual prediction, OR=4.75, 95% CI [2.59, 12.44], whereas the immediate recall condition of VR was the weakest individual predictor, OR=1.91, 95% CI [1.21, 3.43]. However, the strongest prediction came from the latent factor score that incorporated all 7 dependent measures at wave 1, OR=9.88, 95% CI [4.39, 37.72]. Cross-validated receiver operating curves for these three models are displayed in Figure 2a. VR immediate recall had a significantly lower area under the curve (AUC; .570) than CVLT learning trials (.741) or the full latent factor score (.796), both Z>2.65, *p*<.008, but the factor score outperformed the CVLT learning trials at a trend level only, Z=1.81, *p*=.070.

**Table 3.** Odds Ratios and 95% Confidence Intervals from Logistic Regressions of Mild Cognitive Impairment (MCI) Predicted by Baseline Memory Measures

| Dependent Measure | Logical<br>Memory | Visual<br>Reproductions | CVLT | z-score<br>composites | Latent factor<br>scores |
| --- | --- | --- | --- | --- | --- |
| *A. Prediction of Amnestic MCI* |  |  |  |  |  |
| Immediate recall only | **3.01**<br>[1.79, 6.44] | **1.91**<br>[1.21, 3.43] | **3.61**<br>[2.00, 9.39] | **5.27**<br>[2.78, 15.29] | **5.67**<br>[2.92, 16.98] |
| Delayed recall only | **3.22**<br>[1.89, 6.83] | **2.35**<br>[1.50, 4.38] | **3.77**<br>[2.14, 8.48] | **5.86**<br>[3.09, 15.83] | **6.45**<br>[3.30, 17.99] |
| Learning trials only | - | - | **4.75**<br>[2.59, 12.44] | - | - |
| Immediate & delay<br>recall<sup>a</sup> | **3.28**<br>[1.93, 7.05] | **2.40**<br>[1.51, 4.56] | **4.26**<br>[2.29, 10.92] | **6.57**<br>[3.32, 19.91] | **6.91**<br>[3.47, 20.20] |
| Immediate & delay<br>recall + learning trials<sup>a</sup> | - | - | **5.60**<br>[2.83, 16.06] | **8.82**<br>[4.04, 32.61] | **9.88**<br>[4.39, 37.72] |
| *B. Prediction of Non-Amnestic MCI* |  |  |  |  |  |
| Immediate recall only | 1.10<br>[.71, 1.72] | **2.02**<br>[1.33, 3.28] | 1.29<br>[.85, 1.99] | **1.71**<br>[1.08, 2.85] | **1.63**<br>[1.03, 2.69] |
| Delayed recall only | 1.11<br>[.71, 1.77] | **1.55**<br>[1.01, 2.47] | **1.78**<br>[1.15, 3.01] | **1.72**<br>[1.07, 2.96] | **1.86**<br>[1.16, 3.21] |
| Learning trials only | - | - | 1.03<br>[.66, 1.57] | - | - |
| Immediate & delay<br>recall<sup>a</sup> | 1.11<br>[.71, 1.76] | **1.96**<br>[1.27, 3.25] | **1.53**<br>[1.01, 2.47] | **1.78**<br>[1.11, 3.04] | **1.91**<br>[1.19, 3.31] |
| Immediate & delay<br>recall + learning trials<sup>a</sup> | - | - | 1.36<br>[.89, 2.15] | **1.64**<br>[1.02, 2.76] | 1.56<br>[.99, 2.60] |

*Note:* Each cell displays an odds ratio (OR) from a separate analysis in which that memory measure (or combination of measures) predicts progression to amnestic MCI (A) or non-amnestic MCI (B). Significant ORs are displayed in bold ( $p < .05$ ). Like all individual measures, z-score composites and factor scores were scored such that higher ORs indicate greater risk for amnestic MCI (aMCI) or non-amnestic MCI (nMCI) at -1 *SD* for that variable.
<sup>a</sup> indicates measures in this row were also based on z-score composites (e.g., LM immediate and LM delayed recall in the first column), except for the final column which was based on factor scores (from the models displayed in Figure 1c and 1d).

**Figure 2:**
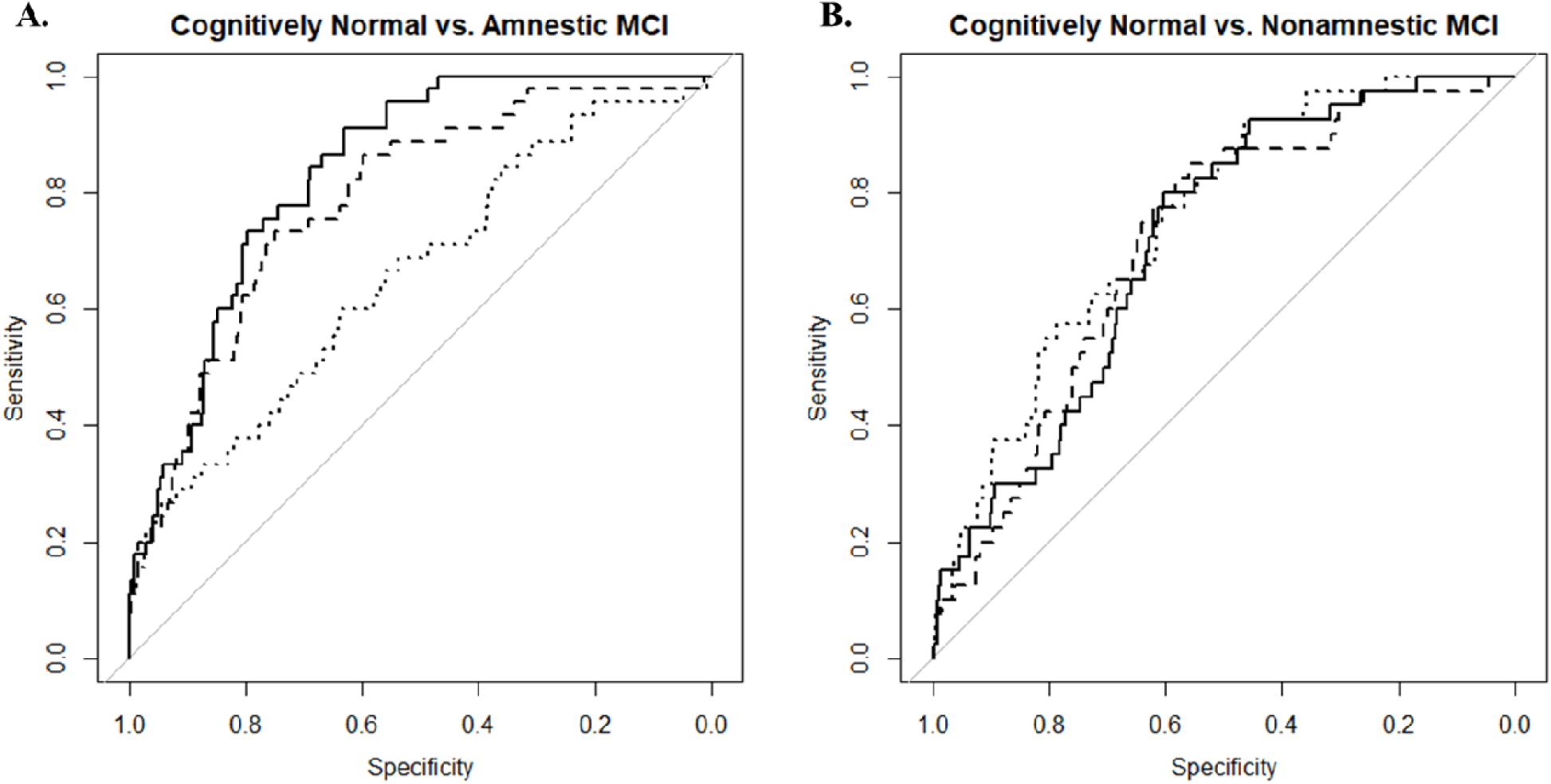
Receiver operating characteristic (ROC) curves for the logistic regressions displayed in Table 3 for mild cognitive impairment (MCI) at wave 2 predicted by memory measures at wave 1. Three models are displayed here: the worst individual predictor of amnestic MCI (VR immediate recall; dotted line), the best individual predictor of amnestic MCI (CVLT learning trials; dashed line), and the best overall predictor of amnestic MCI (latent factor score of all measures; black line). Area under the curve (AUC) estimates were obtained by doing 4-fold cross validation repeated 10 times. Delong’s tests revealed significant AUC differences for amnestic MCI between VR immediate Recall (AUC=.570) and both CVLT learning trails (AUC=.741) and the full latent factor score (AUC=.796), both Z>2.65, *p*<.008, but not between CVLT-Learning Trails and the latent factor score, Z=1.81, *p*=.070. There were no significant AUC differences for nonamnestic MCI, Zs<1.36, *p*>.175 (AUCs for VR immediate recall=.686, CVLT learning trials=.607, latent factor score=.637). VR = Visual Reproductions; CVLT = California Verbal Learning Test.

Although most confidence intervals overlapped, long delay recall conditions tended to be stronger predictors of progression to MCI than short delay conditions (average increase=11.5%, range=4.4%-23.0%). OR estimates for composites greatly outperformed the individual measures. Aggregating measures within the same test resulted in a small increase in ORs (average increase=21.2%, range=2.1%-55.1%). Aggregating measures of the same type across tests resulted in a larger increase in ORs (average increase=105.9% range=46.0%-296.9%). Aggregating across all 7 measures resulted in the largest increase in ORs compared to individual measures (average increase=213.1%, range=85.6%-517.2%). Finally, ORs for latent factors scores were always larger than *z*-score composites (average increase=8.7%, range=5.1%-12.0%).

A different pattern of results was observed for non-amnestic MCI. Some memory measures at wave 1 predicted progression to non-amnestic MCI at wave 2 (3 out of 7 measures), the strongest of which was VR immediate recall, OR=2.02, 95% CI [1.33, 3.28]. However, the predictive ability was weaker than it was for amnestic MCI. Although aggregating measures sometimes resulted in larger ORs than those for individual measures, none of these estimates was stronger than that for VR immediate recall alone. Thus, there was some evidence that baseline memory predicted later non-amnestic impairment, but combining a mix of informative and non-informative predictors did not appear to strengthen prediction.

## Discussion

Here we showed that memory performance among cognitively normal adults who were only in their 50s predicts 6-year progression to amnestic MCI. The results confirmed our primary hypothesis that combining multiple measures of memory improved prediction of progression to amnestic MCI. These results have strong implications for prospective studies aimed at early identification of individuals at greatest risk for MCI and AD. They suggest that longitudinal prediction of amnestic MCI can be substantially improved by both administering multiple memory tests and utilizing multiple scores that are available within each test. The use of a single best measure may appear to be time- and cost-effective. However, in this study, administering neuropsychological tests of both verbal and non-verbal memory added only about 30 minutes of additional test administration time and resulted in a dramatic increase in the odds ratio. Even when more than one test has been used, many studies seldom include more than one score from each test. Calculating additional scores from already completed tests adds nothing to study participant burden.

Predictors of 3-year progression to MCI or dementia in cognitively normal adults in the Australian Imaging, Biomarkers, and Lifestyle Study of Ageing (AIBL) serve as a sample comparison[42]. In AIBL (mean age=72.0 years), Aβ positivity was the best individual predictor (OR=4.8), similar to our CVLT trials 1-5 (OR=4.75). AIBL examined CVLT delayed recall (OR=4.2), which was similar to that demonstrated in VETSA here (OR=3.77). The best prediction in AIBL came from combining Aβ and CVLT delayed (OR=15.9) compared with the prediction based on all memory measures in VETSA (OR=9.88). A total of 13% of AIBL progressed to MCI or dementia whereas 5% of VETSA participants progressed to amnestic MCI. However, the memory tests in VETSA are accounting for substantially earlier prediction because the average age was 16 years younger than in AIBL. Although measures of Aβ may have increased prediction of conversion to MCI in VETSA, we found that cognitive performance alone has good predictive utility. Obtaining biomarkers of amyloid accumulation in a sample with such a low number of individuals who are expected to be Aβ positive [43] would be highly cost-ineffective. Non-invasive neuropsychological tests are cost-effective and sensitive measures for very early risk identification. Using cognitive testing as a first-line screening in clinical trials to reduce unnecessary numbers of PET scans and CSF draws could save millions of dollars and reduce participant burden associated with obtaining biomarker measures of amyloid[44]. Moreover, being Aβ-negative at this age does not necessarily mean non-AD-related as there is also evidence that Aβ at subthreshold levels can still have negative effects on cognition[13]. In any case, it will be important for future work on younger samples to evaluate the joint prediction to see if AD biomarkers improve prediction over neuropsychological tests alone in participant as young as those in VETSA.

Including tests in other cognitive domains might also further improve prediction. Some examples include the preclinical Alzheimer cognitive composite (PACC), which used 4 tests common to 3 samples[45], and multiple tests in the Einstein Aging Study[46] and the Rush Memory and Aging Project[47]. However, these indices were created by and for older samples. As in the large majority of studies, their focus was on predicting progression to AD rather than MCI. Baseline ages of participants for these analyses were 16-24 years older than the VETSA baseline age. VETSA was designed to have an extensive and taxing battery in order to capture heterogeneity and to avoid ceiling effects in our much younger participants[27, 48]. We see our approach as complementary. Instead of including 1 score per test as in these other studies, we examine more extensive coverage within specific domains.

In contrast, combining multiple memory predictors did not increase overall prediction for progression to non-amnestic MCI (though all 95% CIs overlapped). The results argue against general cognitive deficit as a predictor of MCI. Rather, there appears to be some specificity of cognitive predictors of MCI. Thus, combining multiple cognitive measures appears to be capable of dramatic improvement in prediction *if they are domain-relevant*.

### Strengths, Limitations, and Future Directions

First, the sample comprised only men, so it will be important to examine whether these findings generalize to women. Second, amnestic MCI diagnoses were not validated with AD biomarkers. On the other hand, these diagnoses at wave 1 were validated with evidence of reduced hippocampal volume[35] and higher AD polygenic risk scores in those with amnestic MCI[36], the latter supporting their being AD-related. Moreover, the 9% of individuals excluded for MCI at wave 1, and 10% converting to MCI at wave 2 corresponds well to recent estimates that about 10% of individuals in their mid-50s are amyloid positive, with another 10% percent becoming amyloid positive by age 65[43]. We do not expect all our MCI subjects to be exactly the same as individuals who are amyloid-positive, but this correspondence at least suggests the plausibility of this being AD-related MCI. Some may also be at subthreshold amyloid levels, and there is growing evidence that subthreshold levels may still be associated with reduced cognitive function[13]. Third, many confidence intervals overlapped, but a power analysis suggested that a latent factor based on many test scores would substantially reduce the number of participants needed for studies. Given the proportion of amnestic MCI in this sample (5.3%), power estimates using Gpower V3.1.9.2 (without covariates) suggest that a study which administered only logical memory delayed recall (OR=3.22) would require 102 subjects to significantly predict progression to amnestic MCI at the .05 level (with 80% power). When using the latent factor based on all 7 memory scores (OR=9.88) the same power analysis indicates that only 35 subjects would be necessary. Thus, we conclude that the small amount of additional non-invasive time in adding two memory tests is a very worthwhile investment.

Fourth, the result that cognitively normal individuals closest to the MCI cutoff were at greatest risk for later MCI might appear to suggest that some prediction could stem from test/retest noise. However, the fact that *z*-scores and factor score approaches (which reduce measurement error) improved prediction argues against this point. Moreover, post-hoc analyses revealed significant prediction by the latent factor score (OR=3.42) even when the amnestic MCI group was reduced to only the 22 (out of 45 total) individuals who also declined by >1 SD on the full memory factor score between waves 1 and 2. Fifth, as in most longitudinal studies, attriters tend to have lower cognitive ability than the returnees. Indeed, the 242 dropouts had significantly lower memory factor scores at wave 1, *p*=.009. Thus, we may have lost some individuals who were at the greatest risk for later memory impairment, but that would suggest that our findings regarding predictive ability are conservative. Finally, fitting the latent factor model and conducting the logistic regression simultaneously might lead to further improvement of factor scores. Indeed, prediction of amnestic MCI by latent factors in the same model resulted in 37%-57% larger odds ratios than the factor scores displayed in Table 3 (as high as 15.54; see supplement Table S2). However, we presented the results of the 2-step procedure here (exporting factor scores, then running a logistic regression) because they are more conservative and are more generalizable in that they can be used by other researchers and/or clinicians to generate risk probabilities in new samples without having to refit a new latent variable model.

### Concluding Remarks

Baseline memory measures can be very useful for early identification as they strongly predict progression to amnestic MCI in middle-age adults. All individuals were cognitively normal at baseline, but individual differences still effectively predicted MCI 6 years later. Prospective studies of MCI designed to identify those at greatest risk should administer multiple memory tests and utilize multiple scores from each test as early as possible to maximize their ability to predict change. The Alzheimer’s Association has projected that diagnosing individuals in the MCI stage, as opposed to the dementia stage, could improve quality of life and massively reduce the financial impact of the disease[49]. Thus, even moderate additional gain in prediction by using additional tests could save large amounts of money when applied to large populations. It is also worth examining whether and when tests in other cognitive domains might further improve prediction. Nevertheless, the results indicate that neuropsychological assessment can be a sensitive predictor of risk for MCI even at an age when few individuals are likely to have become biomarker positive.

## Acknowledgements

The content of this manuscript is solely the responsibility of the authors and does not necessarily represent the official views of the NIA/NIH, or the VA. The U.S. Department of Veterans Affairs has provided financial support for the development and maintenance of the Vietnam Era Twin (VET) Registry. Numerous organizations have provided invaluable assistance in the conduct of the VET Registry, including: Department of Defense; National Personnel Records Center, National Archives and Records Administration; Internal Revenue Service; National Opinion Research Center; National Research Council, National Academy of Sciences; the Institute for Survey Research, Temple University. This material was, in part, the result of work supported with resources of the VA San Diego Center of Excellence for Stress and Mental Health Healthcare System. Most importantly, the authors gratefully acknowledge the continued cooperation and participation of the members of the VET Registry and their families as well as the contributions of many staff members and students.

## Funding

This research was supported by Grants R01 AG050595, R01 AG022381, and R01 AG059329 from the National Institutes of Health.

## Notes

#### Summary of Updates

Revisions were made during peer review. They include some additional analyses, minor revisions to the introduction, and revisions to the discussion/limitations.

