## Supplemental Information for "Extensive memory testing improves prediction of progression to MCI in late middle age"

### Supplemental Method

**Latent factor scores.** Latent factors were exported from structural equation models in MPlus version 7.2[1]. These factor scores were based in part on models reported in earlier work from this sample[2, 3]. Although there were some methodological differences in this previous work (i.e., all memory measures were adjusted for general cognitive ability, the model was based on a full twin analyses rather than phenotypic data only), Kremen et al. (2014) demonstrated that the hierarchical factor model of memory displayed in Figure 1c provided the best fit to the 6 memory measures (3 immediate and 3 delayed recall) compared to other confirmatory models. We confirmed that this model continued to fit the data well on the non-adjusted memory measures (see below), and therefore did not test any alternate models.

The other models displayed in Figure 1 were adapted from that displayed in Figure 1c depending on the measures included. The models from Figure 1a (immediate recall only) and Figure 1b (delayed recall only) are just-identified, so they have perfect model fit. The model displayed in Figure 1d included the CVLT learning trials, so we included this measure as another indicator of the test-specific factor for the CVLT. However, we also used modification indices to identify an additional factor loading from the CVLT learning trials directly to the latent Episodic Memory factor. This factor loading was not predicted *a priori* but was intuitive given that this is was only memory measure that directly assessed encoding. Results of the model were similar without this additional factor loading, but the model displayed in Figure 1d was more parsimonious (based on all model fit indices) and fit significantly better, $\chi2$ (1) = 12.26, $p < .001$. Additionally, the other factor loadings in the model were similar to those in Figure 1c. There were a few cases where individuals were missing memory data for specific measures (see Table 2). In these cases, *z*-scores and factor scores were created based on all available measures.

Model Fit:

Figure 1a & 1b: $\chi^2$ (0) = 0.00, $p$ = .999, RMSEA = .000, CFI = 1.00

Figure 1c: $\chi^2$ (6) = 8.53, $p$ = .202, RMSEA = .018, CFI = .999

Figure 1d: $\chi^2$ (10) = 47.60, $p$ = .000, RMSEA = .054, CFI = .992

*Table S1*

*Descriptive Statistics for Episodic Memory Measures at Baseline Split by MCI Diagnosis at Follow-Up*

| | Cognitively Normal (N=763) | | | | Amnestic MCI (N=45) | | | | Non-amnestic MCI (N=41) | | | |
|---|---|---|---|---|---|---|---|---|---|---|---|---|
| Memory Variable | N | M | SD | Range | N | M | SD | Range | N | M | SD | Range |
| *Logical Memory* | | | | | | | | | | | | |
| Immediate Recall | 761 | 24.63 | 5.70 | 8, 44 | 45 | 20.20 | 5.40 | 8, 31 | 41 | 23.78 | 5.67 | 12, 39 |
| Delayed Recall | 761 | 21.31 | 6.02 | 3, 40 | 45 | 16.58 | 5.83 | 2, 27 | 41 | 20.44 | 5.51 | 6, 32 |
| *Visual Reproductions* | | | | | | | | | | | | |
| Immediate Recall | 763 | 80.84 | 10.77 | 46, 103 | 45 | 74.31 | 14.64 | 39, 100 | 41 | 72.85 | 12.14 | 42, 98 |
| Delayed Recall | 762 | 58.67 | 17.90 | 0, 99 | 45 | 46.07 | 19.91 | 10, 86 | 41 | 50.29 | 18.63 | 13, 90 |
| *California Verbal Learning Test* | | | | | | | | | | | | |
| Learning Trials | 758 | 5.46 | 1.51 | 2, 13 | 45 | 4.84 | 1.15 | 3, 8 | 40 | 5.48 | 1.83 | 2, 11 |
| Short Delay Free Recall | 755 | 9.17 | 2.60 | 2, 16 | 45 | 7.16 | 1.72 | 4, 12 | 40 | 8.43 | 2.90 | 4, 16 |
| Long Delay Free Recall | 754 | 9.72 | 2.64 | 0, 16 | 45 | 7.49 | 2.19 | 1, 14 | 40 | 8.33 | 2.82 | 3, 15 |

*Note:* Descriptive statistics for memory measures at wave 1 are shown broken down by mild cognitive impairment (MCI) diagnosis at wave 2. In all analyses involving MCI, dependent measures were standardized based on the full sample at wave 1 (see Table 2) and reverse scored so that odds ratios reflect increase in risk of MCI at lower levels of cognitive ability.

*Table S2*

*Odds Ratios and 95% Confidence Intervals from Logistic Regressions of Mild Cognitive Impairment (MCI) Predicted by Baseline Memory Latent Factors*

| Dependent Measure | Latent Factor |
|---|---|
| *A. Prediction of Amnestic MCI* | |
| Immediate recall only | 8.92 |
| | [3.46, 23.00] |
| Delayed recall only | 8.43 |
| | [3.81, 18.70] |
| Immediate + Delay | 9.45 |
| | [4.10, 21.89] |
| Learning Trials | - |
| Immediate + Delay + Learning | 15.54 |
| | [5.38, 44.84] |
| *B. Prediction of Non-Amnestic MCI* | |
| Immediate recall only | 2.26 |
| | [1.04, 4.75] |
| Delayed recall only | 2.05 |
| | [1.11, 3.78] |
| Immediate + Delay | 2.24 |
| | [1.16, 4.32] |
| Learning Trials | - |
| Immediate + Delay + Learning | 1.81 |
| | [.94, 3.13] |

*Note:* These results are identical to those displayed in Table 3 of the main text except MCI diagnoses were predicted directly by the memory latent factors rather than latent factor scores. Each cell displays an odds ratio (OR) from a separate analysis in which that memory latent factor predicts progression to amnestic MCI (A) or non-amnestic MCI (B). All ORs are significant ($p < .05$). Latent factors were scored such that higher ORs indicate greater risk for amnestic MCI (aMCI) or non-amnestic MCI (nMCI) at -1 *SD* for that variable. Analyses were conducted in Mplus using the multinomial logistic regression estimator, type=complex (to control for familial clustering), and monte carlo integration.
